# Correlative MS Imaging for *in situ* cryo-ET

**DOI:** 10.1101/2025.09.16.676641

**Authors:** Andre Schwarz, Jakob Meier-Credo, Arne Behrens, Jürgen Reichert, Sonja Welsch, Lea Dietrich, Erin M. Schuman, Julian D. Langer

## Abstract

*In situ* cryo-electron tomography (cryo-ET) offers direct access to protein structures in their native state and environment. A central challenge for this technique is the identification and classification of imaged cells, as sample heterogeneity with respect to cell type and functional state can significantly impact the data. Currently, cells can be classified using subtomogram averaging or fluorescent labeling, which require sufficiently distinguishable macromolecules, genetic modifications, or the availability of suitable labels. In MALDI-MS Imaging (MSI) individual cells can be identified and classified reliably, making use of the distributions of phospholipid species, metabolites and small proteins with characteristic profiles for specific cell types. Here we present an integrated pipeline combining MSI and *in situ* cryo-ET for single-cell identification and classification. Our approach enables MSI data acquisition directly from EM grids and can be used on any eukaryotic single-cell cryo-ET sample without prior modification in less than 36 h. We apply this workflow to describe cell-to-cell heterogeneity in cultured HeLa cells, to distinguish between distantly related HeLa and *Polytomella* cells, and between primary cultured rat neurons and glial cells. Finally we present a data integration workflow combining cryo-ET, light microscopy and MSI aligned in the same coordinate system to facilitate efficient data visualization and interpretation. We anticipate that this approach will change the way that structural biologists can approach cellular heterogeneity in cryo-ET, as cells can be identified and classified reliably post acquisition.

## Introduction

*In situ* cryo-electron tomography (cryo-ET) has revolutionized structural biology: structures of protein complexes are accessible in their native cellular environment both in cultured cells (Mahamid et al., Science, 2016 ) and tissue samples (Schaffer et al., Nature Methods, 2019, Schiøtz et al, 2024; Nguyen et al., 2024). This technique has provided new insights into e.g. the mode of action of clinical drugs (Xue et al., Nature 2022, Xing et al., Science, 2023), muscle fiber organization (Wang et al., 2021), and the first molecular description of new microorganisms (Rodrigues-Oliveira et al., 2023). A key challenge for this method is the unambiguous determination of the identity and state of the cells from which protein structures are obtained. Particularly in environmental samples or primary cell cultures, with multiple organisms or cell types present, the identity of a cell of interest has to be inferred from the cryo-EM data directly (Rodrigues-Oliveira et al., 2023) or via correlative labeling techniques (Pierson et al., 2024; Fung et al., 2023). These methods can, however, affect or impair sample quality, rely on availability of suitable reagents, and often yield limited information. In addition, they all report the distribution of selected markers and, while offering sensitive and robust detection for a specific target, may not capture cellular heterogeneity or different functional states of cells. Thus even for comparatively homogeneous samples, e.g. a cell line in culture, the metabolic state of a cell remains unclear.

MALDI-MS data is routinely used to identify microorganisms in clinical samples. The small molecule, lipid, and small protein signal patterns are highly characteristic and utilized in clinical “Biotyping” to classify medically-relevant pathogens based on their MALDI mass spectra signatures. At present, more than 500 strains of human pathogens can be identified with clinical validation (99.9% ID success rate) and more than 2,000 strains are available in reference databases (with >98.3 ID success rate). MALDI-MS thus represent a powerful tool for tissue and cell identification and characterization.

While Biotyping is conducted on extracts and homogenized samples, MS Imaging (MSI) investigates the spatial distribution of metabolites, lipids, and small proteins in intact cells and tissues. MSI data are usually acquired at micrometer resolution and can therefore be used to differentiate cell types in mixed cultures or tissues, and to characterize and differentiate their metabolic states (Rappez et al., Nature Methods, 2021). A recent study using a combination of immunolabeling-based cell classification and MSI-based lipid profiling enabled the determination of specific lipid patterns for different neuronal and glial cell types (astrocytes and oligodendrocytes, Asadian et al.,2025). Recent studies also used other mass spectrometry-based imaging techniques such as secondary-ion mass spectrometry (SIMS) to directly study cryo-ET samples (Ochner et al., 2025) using a prototype instrument that enables direct SIMS-imaging of cryo-ET grids. This approach offers complementary, submicrometer-resolution spatial information on small, inorganic molecule distributions in the sample. However, biomolecules such as lipids, signaling molecules and peptides are not directly accessible using conventional SIMS.

Here we present a correlative workflow for cryo-ET analysis **(Figure 1A)** utilizing high-resolution MSI to analyze individual cells following structural analysis and determine their lipid and small molecule profiles. This allows for the unambiguous identification and classification of single cells by species and type, and provides molecular information complementing structural studies.

**Figure 1:**
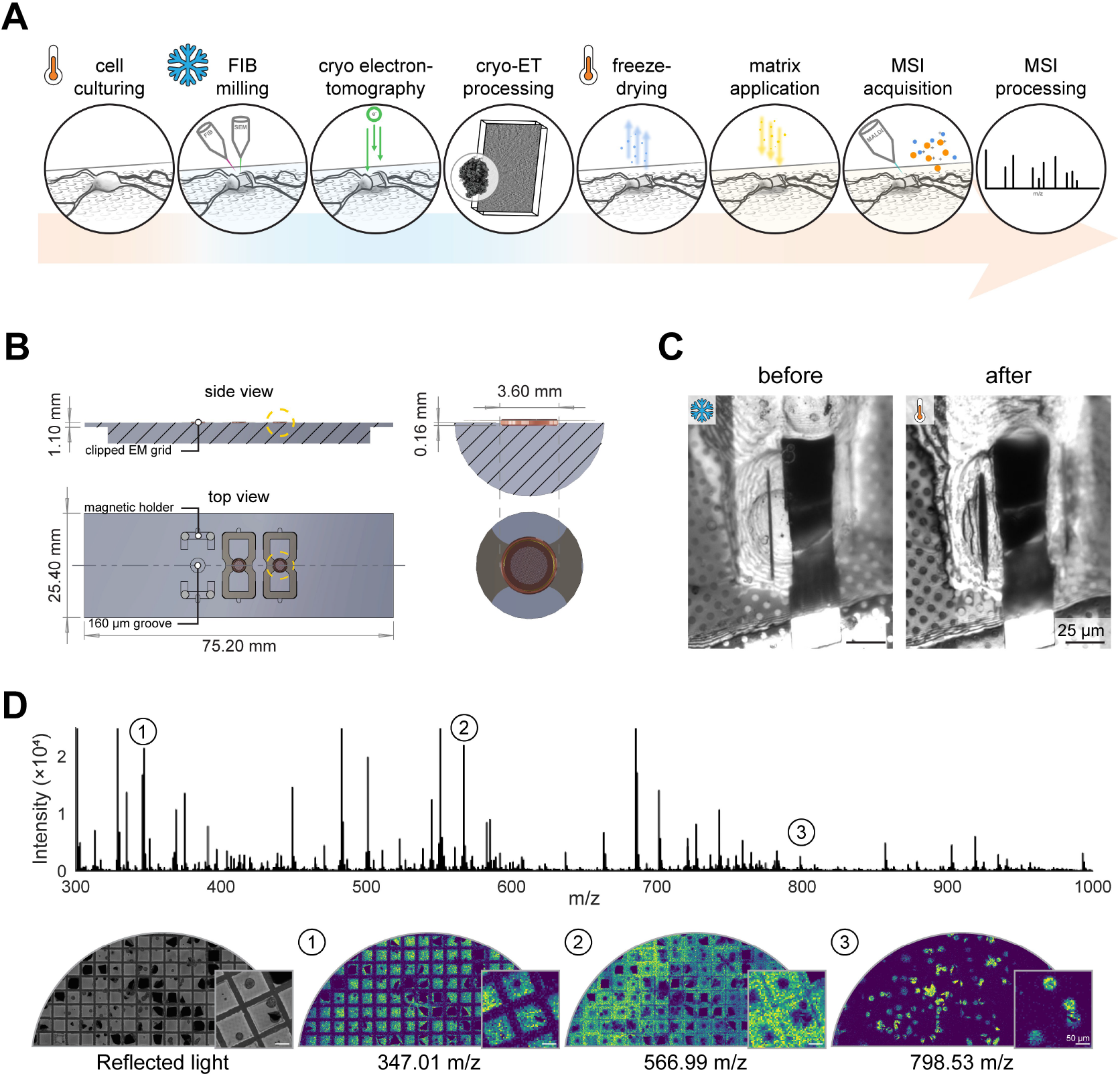
Correlative cellular cryo-ET and MSI enables molecular profiling on plunge-frozen grids. **(A)** Schematic representation of correlative cellular cryo-ET and MSI workflow. **(B)** Technical drawing of custom built grid holder for MSI of clipped cryo-EM grids. Grids are placed within a 160 µm indentation and held by metal springs via a magnetic holder to facilitate an even imaging surface. **(C)** Reflected light images of a HeLa lamella before and after freeze-drying in a dual beam microscope column. Gradual water sublimation leads to cell shrinkage visible by changing interference patterns. **(D**) Example average mass spectrum, individual mass to charge (m/z) channels, and reflected light images of an EM grid containing HeLa and *Polytomella* cells. Different m/z channels display specificity to background support, uniform signal across the grid, and cellspecific signal respectively (from left to right). Y-axis is capped at 2.5 × 10^4^ for improved visibility of smaller peaks.

## Results

### Adapter and workflow

In order to enable a streamlined transition of samples between EM and MSI, we first set out to construct an adapter that enables MALDI-based acquisition directly on EM grids **(Figure 1A)**. Grids for cryo-ET are typically “clipped” into small holder rings for transfer within and between electron microscopes, so the plane of the grid (and cells) is approximately 160 µm above the bottom edge of the clip rings. In order to secure grids into regular cover slide MSI adapters, we designed and produced corrosion-resistant aluminum grid holder adapters using CAD and high-precision milling. Due to spatial limitations of the laser optics and instrument ion lenses, the adapter houses the EM grid in a recess secured by flat springs attached to embedded magnets in the adapter base **(Figure 1B)**. Next, we needed a gentle way to transfer samples from a frozen-hydrated state back to room temperature for MSI pre-processing. Conventional drying approaches were inconsistent and led to cell rupturing and a loss of high resolution spatial information. We thus optimized gentle freeze-drying directly in the electron microscope **(Figure 1C, Figure S1, Movie S1, see methods)**, without any washing steps.

To obtain data with high spatial resolution (at a pixel size of 5 x 5 µm), we finally applied 2,5-Dihydroxybenzoic acid (DHB) matrix using a sublimation workflow. We then tested our adapters on Bruker rapifleX and timsTOF fleX instruments. We obtained robust and sensitive signals on both instruments, but due to the clip ring geometry and the instrument laser angle, we were only able to achieve complete grid coverage on the timsTOF fleX. When combined, this approach yielded grids with structures matching cell body positions prior to drying, with superior cellular anatomy and integrity **(Figure 1D)**. In this pilot study, we focused on lipid profiling as these molecules are highly abundant, typically display high ionization efficiencies in MSI and also provide characteristic signal patterns for different cell types (Rappez et al., Nature Methods, 2021). We observed clear signals in the typical phospholipid range at these cell body locations. Post drying, MSI data for a complete grid was acquired within 12 h at 5 µm pixel size. Notably, some of the cell preparations used here were stored for up to several years in liquid nitrogen prior to freeze-drying, and then several months in a desiccated environment between freeze-drying and MSI analysis. No significant additional damage was observed to the carbon film on the grids during freeze-drying.

### Cell identification and characterization: Cultured HeLa cells

Having established a robust sample preparation pipeline, we next tested our approach on different model systems. First, we investigated if we could identify and characterize individual HeLa cells grown on EM grids. Notably, these cells were vitrified directly in medium without any washing steps to maintain fully physiological conditions. Consequently automated peak picking identified thousands of m/z peaks **(Figure 1D)**, with the majority of these signals deriving from components of the medium, the grid support and the carbon film. In order to comprehensively identify cell-specific peaks, we first selected a set of peaks matching previously identified phospholipid components of the HeLa lipidome such as PC 34:1, PC 36:2 and PE 36:1 and found that their locations matched cellular locations from optical images **(Figure S2)**. We then further annotated 73 phospholipid species in database searches of different HeLa grid preparations **(Table S1)**, conducting on-grid MS/MS measurements of selected candidates to confirm the database searches **(Figure S3A-B)**. We observed differences in both cell size, and shape, as well as levels of cation adduct formation between different preparations and cell batches. However, overall, we identified identical phospholipid species across all experiments with similar intensity profiles.

The identified peaks then served as markers in a comprehensive all-vs-all spatial cross-correlation analysis and subsequent hierarchical clustering **(Figure 2A, see methods)**. As expected, most of the clusters corresponded either to “background” channels covering grid bars and/or the carbon support, or noise channels without specific spatial pattern. Two clusters, comprising a total of 97 m/z channels however contained both the majority of marker channels and were specific to cellular locations, as judged by correlated reflected light images **(Figure 2A, Table S2, Figure S4)**. We thus conclude that we can reliably and robustly detect and identify HeLa cells on EM support grids, and comprehensively identify cell-specific m/z channels via spatial cross-correlation.

**Figure 2:**
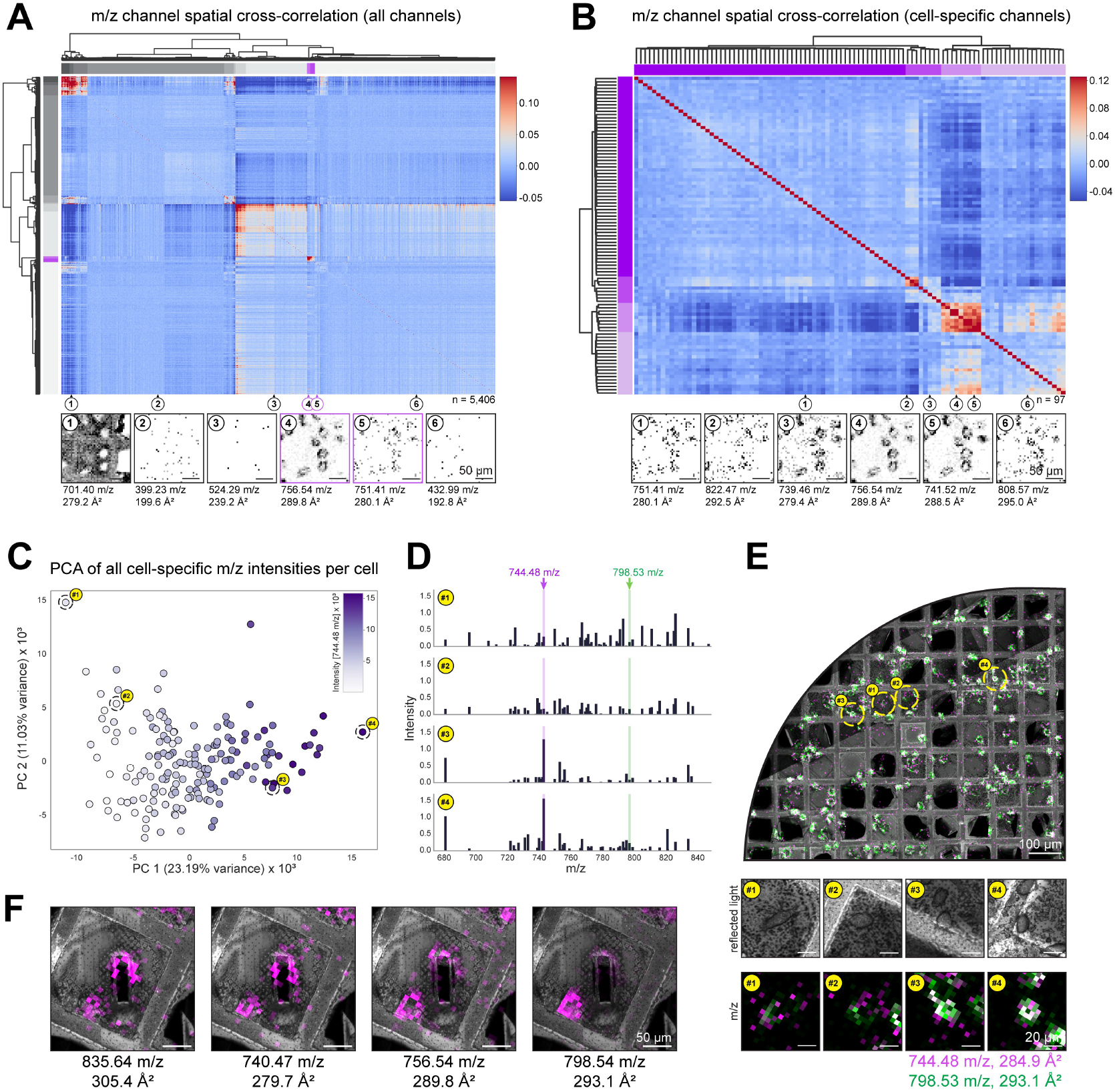
Mass spectrometry imaging of HeLa cells and lamellae shows significant variability between cells. **(A)** Hierarchical clustering of normalized spatial correlations to identify cell-specific m/z channels (n = 5,406). Example m/z channel images of selected clusters show prominent features of each cluster. Cell-specific clusters are highlighted in purple. **(B)** Second round of hierarchical clustering of only cell specific channels from (A) (n = 97). Different clusters largely display different signal-to-noise ratios and/or heterogeneity between cells. **(C)** Principal component analysis of all individual cells over all cell-specific m/z channels in (B), colored by the intensity of m/z 744.48 (PC 30:0). A continuous distribution is observed with 744.48 m/z accounting for 65% of PC1. Individual cells along PC1 are highlighted by yellow circles (containing cell ID’s). **(D)** Spectra for highlighted cells in (C) covering cell-specific m/z values from (B). X-axis is cropped to a lipid-rich region between 675 - 850. Shaded areas highlight variable m/z channel 744.48 (magenta) and stable channel 798.53 m/z (PC 34:1, green) displayed in (E). **(E)** Reflected light overview of HeLa cells grown on an EM grid with highlighted m/z channels from (D) overlaid. Individual cells from (C, D) are highlighted and enlarged. m/z channel 744.48 intensity (magenta) increases from cell #1 - #4. **(F)** Reflected light images and m/z channel overlays of channels showing enriched signal at regions adjacent to lamellae. Adjacent regions of cut lamellae retain clear MSI signal.

Next, in order to investigate inter-cellular variability within this dataset, and whether we could discern distinct functional states among cultured HeLa cells, we performed a second round of hierarchical clustering of these cell-specific m/z channels **(Figure 2B)**. This yielded a cluster of m/z channels with strong correlation and signal-to-noise-ratio, alongside several clusters of more variable channels. By correlating a reflected light-based mask for each cell **(Figure S4A)**, we then measured the mean intensity for each of these 97 cell-specific m/z channels per cell. When viewed as a principle component analysis (PCA) of all cells across all m/z channels, a continuum emerged with cells at opposite ends exhibiting considerable variability for different m/z values **(Figure 2C)**. The most variable channel, PC 30:0 (744.48 m/z, 284.9 Å^2^), mirrored the largest axis of this variability along PC 1. Consequently, cells from opposite ends showed a strong difference in lipid content. When displayed alongside a more stable channel, PC 34:1 (798.53 m/z, 293.1 Å^2^), the increase in PC 30:0 was clearly noticeable in individual cells **(Figure 2D-E)**. This cellular heterogeneity is directly in line with previous MSI studies on HeLa (Xi et al., 2020; Otsuka et al., 2025) and other cell lines in culture (Bien et al., 2022), showing cell-specific, distinct phospholipid profiles with varying lipid compositions.

The analyses thus far were based on intact HeLa cells. In a standard cryo-ET workflow of cultured cells, however, “lamellae” are cut from target cells using a focused ion beam (FIB). This process, called “FIB-milling”, generates electron-translucent thin sections of the target cells for high-resolution structural analysis, at the cost of removing a large fraction of their volume. We thus examined whether sufficient m/z signal was still detectable on or surrounding these lamellae. For assignment and coregistration of MSI with other imaging modalities, we made use of the regularly spaced grid pattern of the underlying electron microscopy supports, which is clearly visible in all electron-, photon-, and some m/z channels **(Figure 1D, 2E)**. This underlying data structure enabled us to achieve high-accuracy co-registration of cryo-ET, optical, and MSI data, even though the data is acquired at very different pixel sizes. This co-registration using a high number of evenly spaced reference points further allowed us to get a spatial estimate of the registration precision across the grid, ranging from 1.07 to 2.34 µm **(Figure S4B)** – less than half a pixel in MSI. After registration, we observed that regions containing lamellae were clearly visible both in the reflected light and the MSI data **(Figure 2F)**. The remaining cellular structures surrounding the lamellae still yielded strong and reliable signals, clearly distinguishable from the background. These peaks include the main phospholipid species detected in unmilled cells, with similar intensities and distributions, and clearly differ from the surrounding carbon film. These data thus demonstrate that we can identify milled cells and obtain quantitative information on their lipid composition, despite FIB-milling.

### Assignment of cell species: HeLa-*Polytomella* “species-mixing” experiment

We next investigated if the data we obtain in such an experiment are sufficient to identify and classify cells from different organisms. To this end, we conducted a “species-mixing” experiment using HeLa and *Polytomella* cells. *Polytomella* is a non-photosynthetic green alga which is significantly smaller than HeLa cells, but forms aggregates in solution leading to a wide range of clusters of varying size between 5 and 30 µm, and are thus difficult to distinguish from single HeLa cells. We prepared a set of grids in our cryo-ET workflow which contained only *Polytomella* cells, only HeLa cells, or a mixture of both. We first examined the molecular distributions on our *Polytomella*-only grids and, using the previously described spatial cross-correlation analysis, found specific peaks matching the localization of *Polytomella* cells or clusters based on optical images. These peaks mainly corresponded to different phospholipid species, which we matched to lipids detected in screening LC-MS lipidomics runs of *Polytomella* samples (list of matched lipids is available in **Tables S3-4**). We also conducted on-grid MS/MS measurements of selected candidates and confirmed their identifications using the respective fragment ion spectra **(Figure S3C-D)**. Taken together, we observed characteristic phospholipid species for *Polytomella* cells and cell clusters in addition to our list of previously determined HeLa-specific phospholipids.

We then analyzed the grid containing a mixture of HeLa and *Polytomella* cells. As before, we conducted spatial cross-correlation and hierarchical clustering analysis on the full MSI dataset and found a total of 89 peaks with cellular localization patterns, forming two broad clusters **(Figure 3A, Figure S5, Table S5)**.

**Figure 3:**
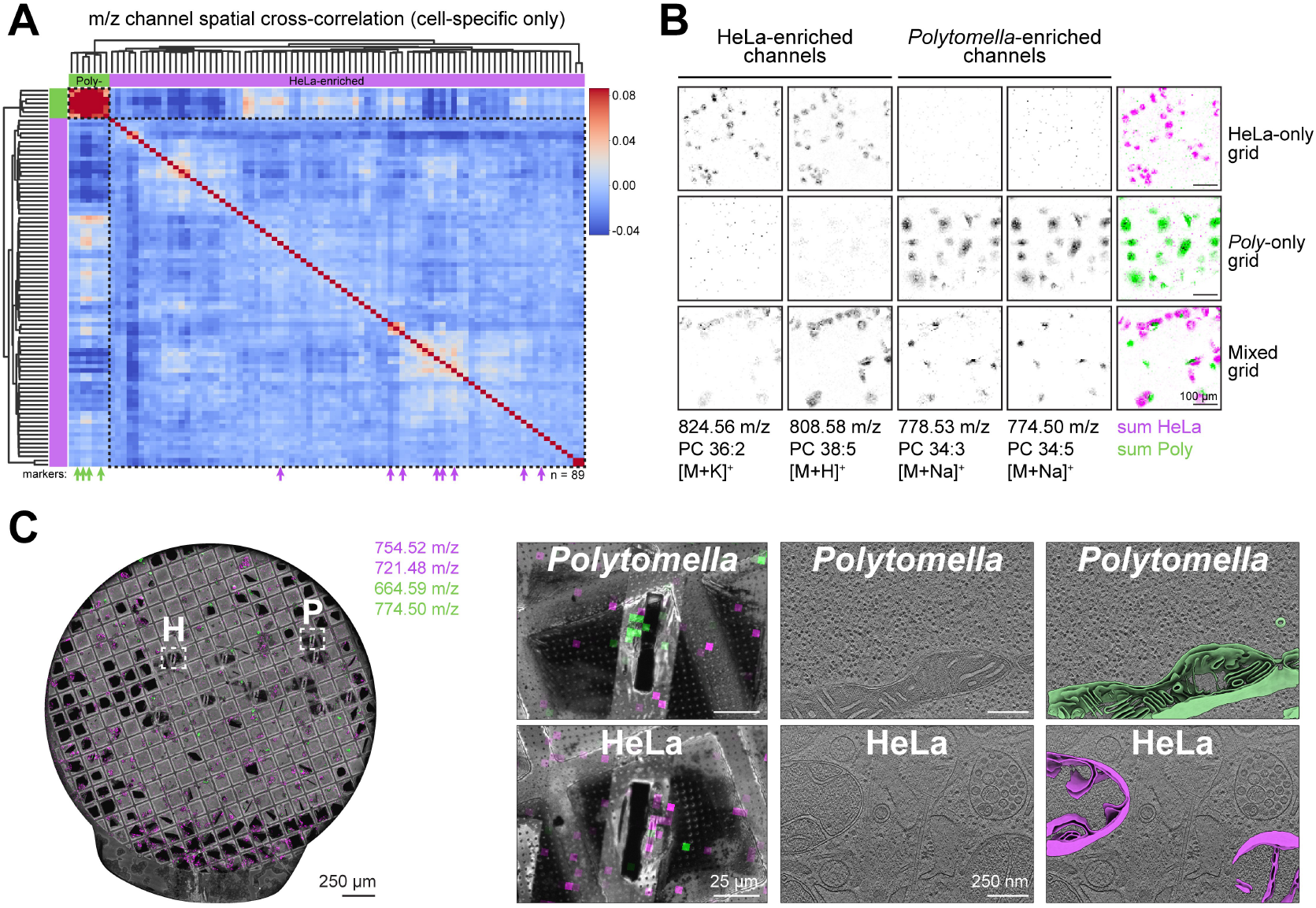
Mass spectrometry imaging signal can distinguish different species in mixed cryo-ET samples. **(A)** Hierarchical clustering of cell specific m/z channels from mixed HeLa / *Polytomella* grid (n=89) with two prominent clusters. Marker channels identified on HeLa-only and *Polytomella*-only grids (magenta and green arrows) identify the clusters as *Polytomella*-enriched and HeLa-enriched m/z channels respectively. **(B)** Example m/z images from both clusters across HeLa-only, *Polytomella*-only, and mixed grids. Both sets of channels show clear spatial separation. **(C)** Reflected light overview of mixed HeLa / *Polytomella* grid with overlaid specific m/z channels from (A). Highlighted regions indicate H = HeLa, P = *Polytomella*. Enlarged images of two lamellae with enriched *Polytomella*- or HeLa-enriched m/z channels with corresponding slices of tomograms and segmentations acquired from these lamellae. Distinct and characteristic mitochondrial shapes confirm the molecular identification.

The individual marker channels identified on *Polytomella*-only and HeLa-only grids served to identify the clusters as *Polytomella*-enriched and HeLa-enriched m/z channels respectively **(Figure 3A, green and magenta arrows)**. These signature channels comprise mainly phospholipids such as PC 36:2, PC 38:5 (HeLa) and PC 34:3, PC 34:5 (*Polytomella*), which were highly enriched on their respective grids and spatially segregated on the mixed grid **(Figure 3B)**.

Making use of these discriminatory m/z channels, we explored whether the remaining MSI signal of FIB-milled cells was sufficient for unambiguous assignment of HeLa and *Polytomella* cells on EM grids. Indeed, based on the signature m/z channels identified in our unsupervised clustering, we were able to unambiguously assign a species identity for 7 out of 10 lamellae cut on the mixed grid **(Figure 3C)**. This classification was possible even though the lamellae were cut from the center of the cells, and the remaining densities often comprised less than 20 MSI pixels. It is worth noting that MSI intensities were generally lower in areas where cells were viewed and cut extensively with the scanning EM (SEM) and FIB beams, likely due to radiation damage. Hence some cells did not have enough signal left for unambiguous identification (which was the case for the remaining 3 out of 10 lamellae).

In order to evaluate our MSI-based classification, we next examined the cryo-ET data recorded from these lamellae. Serendipitously, the tomograms often allow assignment of the two cell types as *Polytomella* cells contain mitochondria with highly stereotypic, disc-shaped cristae, clearly distinguishable from the tighter and less regular cristae in HeLa cells (Dietrich et al., 2024, Medina et al., 2025). The cryo-ET-based classification matched the MSI-based classification for all lamellae for which we obtained tomograms containing mitochondria **(Figure 3C)**. This demonstrates that MSI can indeed be used to identify and classify single cells of distantly related species on heterogeneous EM grids after cryo-ET sample preparation and data acquisition.

### Classification of cell types: Primary neuron cultures

Next we examined primary cultured rat hippocampal cells grown on EM grids that had previously been acquired (Schwarz, Mueller et al., in revision) and remained in liquid nitrogen storage for over two years between tilt series and MSI data acquisitions. These preparations represent a challenging sample as they form dense networks with extensive processes (axons and dendrites; collectively called “neurites”) and contain multiple cell types (neurons and glia cells). Prior fluorescence imaging using NeuO, a neuron-specific fluorescent dye (Cheng Er et. al, 2015), enabled us to identify neuronal positions on the EM grid and match structures visible in reflected light images **(Figure 4A)**. We found clearly distinguishable phospholipid peaks, with multiple phospholipids assigned using high-resolution MS and ion mobility data **(Table S6)**. These lipid species clearly matched not only the locations of cell bodies, but also followed the growth pattern of even individual thin neurites along the grid **(Figure 4A)**. We again confirmed our database matches by acquiring MS/MS data on selected candidates **(Figure S3E-F)**.

**Figure 4:**
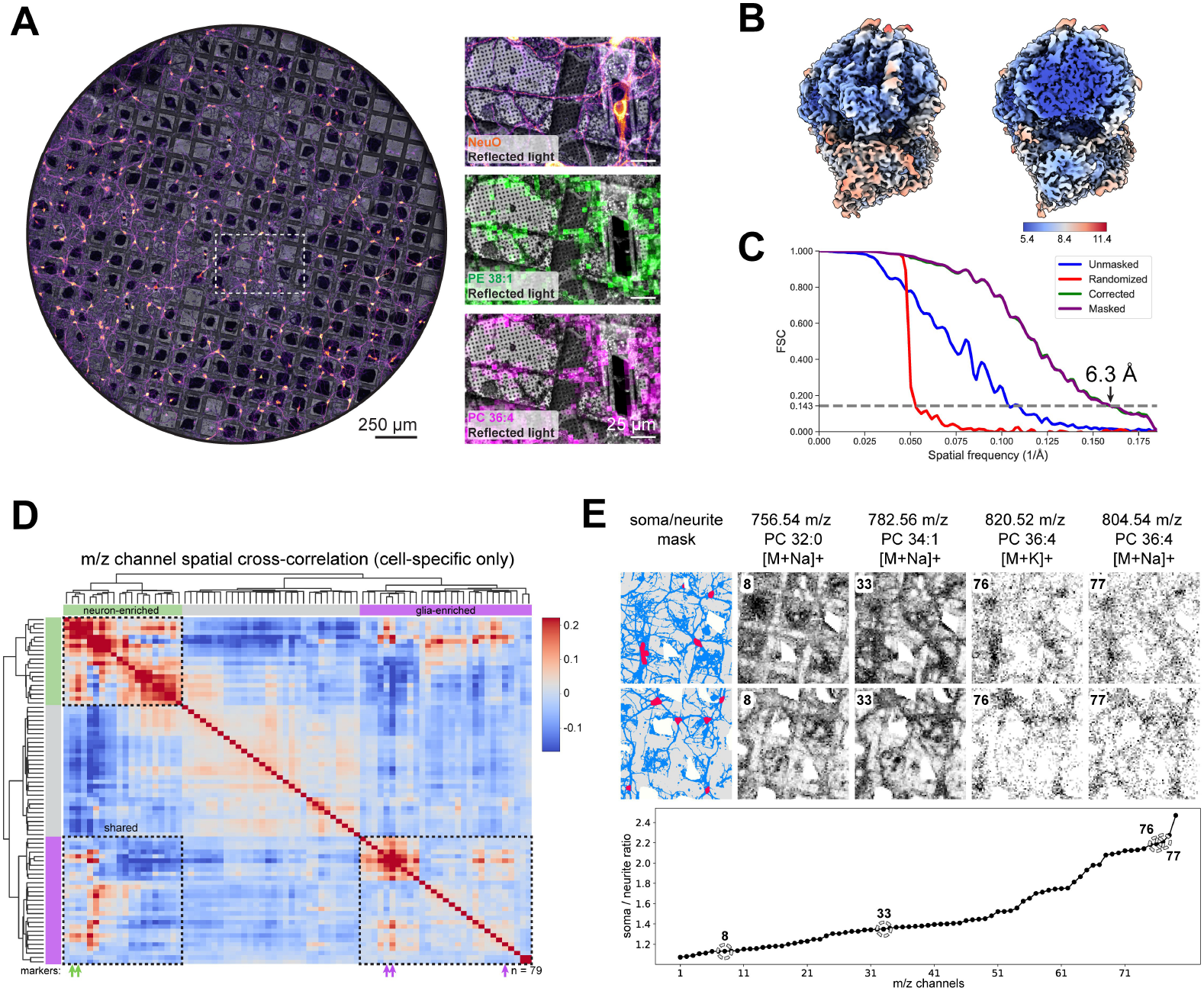
Mass spectrometry imaging of primary neuron culture grids. **(A)** Reflected light overview of primary neuron grid with correlated neuron-specific NeuO live-cell dye and cell-specific m/z channels. **(B)** Subtomogram average (STA) of 15,451 80S ribosomes, extracted from the grid in (A), color-coded by local resolution (in Å). **(C)** Fourier shell correlation plot of STA in (B) with global resolution of 6.3 Å at 0.143 criterion. **(D)** Hierarchical clustering of cell specific m/z channels from primary neuron grid in (A) (n=79) with three prominent clusters. Marker channels specific for neurons (green arrows) or glia cells (magenta arrows) identified in (Asadian et al, 2025) identify the clusters as neuron-enriched and glia-enriched m/z channels respectively with elevated cross-correlation between many channels (“shared”). **(E)** m/z intensity ratios between neuron soma and neurite regions. Individual m/z channels with high or low soma/neurite ratio are highlighted and representative images are displayed alongside the mask used for quantification. Blue = neurites, red = somata.

We then investigated whether we could obtain data from cells that were subjected to FIB-milling. As before, lamellae were cut from the cells of interest in such a way that the majority of the cell body was removed **(Figure 4A)**. Nevertheless, we were able to record strong cell-specific m/z signal surrounding (and sometimes on) lamellae, although it was more difficult to assign this intensity to a specific cell owing to the highly intertwined network of overlapping neuronal structures **(Figure S6A)**. We further noticed that the areas directly surrounding these milled segments displayed characteristic changes, with a moderate signal reduction in some channels, and a distinct signal increase in others **(Figure S6B)**. We speculate that these changes could be due to radiation damage during the FIB-milling process as noted for HeLa/*Polytomella* before, however, cellular analysis was not compromised as the affected channels were not cell-specific. As a proof of concept for the full workflow, we extracted and analyzed tilt series acquired only from this grid’s lamellae, constituting a subset of our previous dataset (Schwarz, Mueller et al., in revision). This resulted in a 6.3 Å global resolution subtomogram average of ∼15,000 rat neuronal ribosomes **(Figure 4B-C**). This shows that the *post hoc* cellular MSI analysis introduced throughout this work can be performed on grids yielding state-of-the art structural data.

Primary brain tissue preparations contain multiple cell types such as neurons and glia. We thus investigated if we could identify and discern these different cell populations in our MSI data. For this, we once again conducted unsupervised hierarchical spatial correlation clustering **(Figure S6C)**, resulting in an early split of three prominent clusters **(Figure 4D)**. In order to test whether any of these corresponded to specific cell types, we used a recently published fluorescence-guided single-cell MSI dataset matching lipids to individual brain-derived cell types via antibody staining (Asadian et al., 2025) as ground truth. When we queried neuron-enriched vs glia-enriched m/z channels **(Figure S6D)**, we indeed found that they separated into two of the three clusters **(Figure 4D, green and magenta arrows)**. Notably, many of these phospholipid species were not unique but merely enriched, with high levels of correlation between them **(Figure 4D, “shared”)**, as observed by others (Asadian et al., 2025).

Having established that we can distinguish neuron- vs glia-enriched m/z profiles on our mixed primary hippocampal culture grids, we next queried whether any of these cell-specific channels displayed any preferential spatial partitioning between cell bodies and neurites. In order to test this, we utilized the previously acquired fluorescence microscopy images of NeuO to create masks for neuronal somata and neurites **(Figure S6E, see methods)** and registered them to the MSI channels with a target registration error to of 0.9 - 2.5 µm **(Figure S6F)**. We then measured the mean intensity for each cell-specific m/z channel for each of these two compartments. We noticed a wide range of partitioning, ranging from channels with a ∼2.5x enrichment in somata (e.g. PC 36:4 (804.54 m/z), **Figure 4E**) to roughly even distributions (e.g. PC 34:3 (756.54 m/z), **Figure 4E**). We conclude that we can still detect lipid distributions in neuronal processes even though their volumes and associated signal intensities are significantly lower than for somata, and that many lipid species display distinct subcellular distributions in primary cultured neurons (Castro et al., Analytical Chemistry, 2023) .

### Interactive data integration in *SpatialData*

Finally, in order to enable streamlined analysis of correlated reflected and fluorescence light, cryo-ET and MSI data, we set out to integrate the different imaging modalities in a single software environment. Making use of the *SpatialData* framework (Marconato et al, 2025) **(Figure 5A-B)**, we constructed an integrated pipeline for registration of all imaging modalities, as well as conversion scripts and notebooks to stitch, visualize and analyze them together. Besides these primary data modalities, derived data such as masks, reconstructed tomograms, segmentations, and particle coordinates including macromolecular activity states can also be integrated and queried without additional effort **(Figure 5B-C)**. Fluorescence and MSI data can thus be directly linked to individual lamellae, enabling rapid, comprehensive analysis of that cell’s molecular contents and ultrastructure.

**Figure 5:**
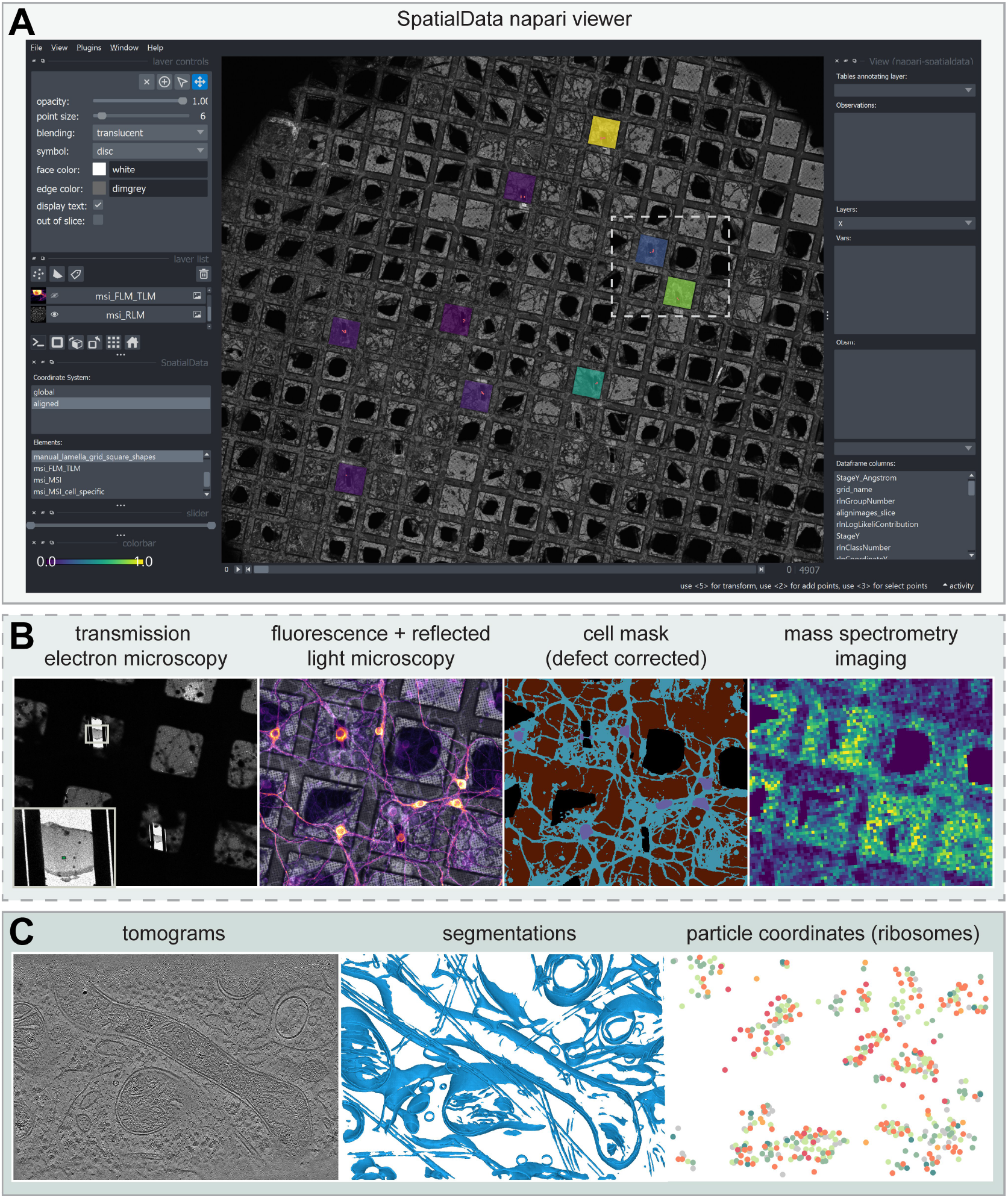
Multimodal integration of MSI and optical microscopy data in *SpatialData*. **(A)** *SpatialData* napari viewer interface with displayed primary neuron grid reflected light image and lamella grid square shapes. Grid square shapes are color-coded by percentage of all active ribosomes (viridis) within each lamella as determined by subtomogram averaging and classification. Values are attached to each lamella shape in the *SpatialData* object. **(B)** Enlarged views of area highlighted in (A) for integrated transmission electron microscopy, reflected and fluorescence light microscopy, cell masks, and a single mass spectrometry imaging m/z channel (from left to right). **(C)** Enlarged views of lamella area highlighted in (B, left panel inset) showing a single tomogram slice, membrane segmentation of the corresponding tomogram, and particle coordinates for identified ribosomes as points (from left to right). Ribosome points are color-coded by translation state from active (greens) to inactive (reds).

## Discussion

Here we present a novel integrated pipeline combining MSI and *in situ* cryo-ET. We developed a workflow that enables MSI data acquisition directly from EM grids that were previously analyzed using FIB-milling and cryo-ET. We first applied our pipeline to HeLa cells and demonstrated sufficient resolution and sensitivity to detect individual cells and describe the intercellular variability in lipidome composition. Our single-cell HeLa data match the results of other groups studying HeLa cells using MSI, with similar lipidome compositions and heterogeneity (Schober et al., 2012; Xi et al., 2020; Otsuka et al., 2025). Examining FIB-milled sections, we observed clear signals for cell bodies with the signature m/z channels of uncut cells. We next conducted a species-mixing experiment to show that we can differentiate between HeLa and *Polytomella* cells — two distantly related species. Using unsupervised clustering, we found that the MSI data segregates clearly into two groups, allowing unambiguous assignment of the two species. Next we examined if we could apply our workflow to primary cultured brain cells, which contain multiple cell types and grow much more densely. Unsupervised clustering of the MSI data allowed clear assignment of glia cells and neurons, making use of previously published data on cell typespecific lipids (Asadian et al., 2025) as marker channels. We also found that certain phospholipids were more abundant in the soma compared to neuronal processes, highlighting the potential of the technique to study sub-cellular lipidome composition and dynamics.

From a technical point of view, there were two main challenges to overcome: A gentle return of frozen-hydrated cryo-ET samples back to room temperature, and the direct loading of cryo-ET grids into the mass spectrometer. Freeze-drying directly in the stage of the electron microscope proved to be a gentle and direct way to preserve cellular structures without damaging the grid. Other drying approaches led to cell rupture, inducing a loss of spatial information and signal intensity. Notably, in this approach, we retain all components of the medium the cells were grown in, as no additional washing steps are conducted. Nevertheless, cell-specific m/z peaks were clearly separable from grid-wide signal. Initial attempts of the full drying process in the cryo-stage took around 24 h but were later routinely carried out remotely overnight. Even though this step does not require any hands-on effort, it is currently the most time-consuming part of the workflow but could certainly be optimized in the future. For straightforward grid transfer, we designed a holder that fits directly into standard MALDI adapters and can also be used for matrix application (e.g. by sublimation). Corresponding CAD files to reproduce this holder are publicly available (see methods). To secure the grid tightly while ensuring accessibility of the laser, the use of embedded magnet-held flat springs was essential, as pressure changes in the instrument could otherwise push the EM grids off the holder and into the instrument.

Our approach addresses one of the key challenges for *in situ* cryo-ET: establishing or confirming the identity of FIB-milled cells, and differentiating their cell type and possibly even metabolic state. At present, cellular identity and state can typically be determined using two main approaches: 1) Correlative imaging via expression of fluorescent (or otherwise recognizable) proteins in the target cells, or application of specific dyes (Pierson et al, 2024; Fung et al, 2023) including fluorescent indicators (Jung et al, 2025). 2) Subtomogram averaging of corresponding macromolecules that are informative about the cell identity or state (e.g. ribosomes) (Rodrigues-Oliveira et al, 2023). Fluorescence imaging and the use of specific dyes offer high confidence cell identification prior to target selection, but require either genetic modification of the target cells or rely on the availability of such dyes. They also might lead to changes in the target cell’s metabolism and structure. In addition, targeted detection of a single reporter protein may not capture the full heterogeneity of a sample. The use of subtomogram averaging and classification of relevant macromolecules on the other hand allows insights on cell identity or state from within the same dataset, but it remains time consuming and requires extensive knowledge of subtomogram averaging and the target cell’s biology. It further relies on the fact that these macromolecules can be identified, averaged, and are sufficiently abundant, informative, and distinct. Compared to these approaches, MSI combines the advantages of not requiring any sample modifications prior to analysis, with fast data acquisition in less than 36 h after cryo-ET acquisition (for a full EM grid at 5 µm pixel size: <24 h freeze-drying, 2 h sample processing, 10 h data acquisition), and provides comprehensive data of the cellular lipidome and other small molecules. Notably, the time for data acquisition could be reduced significantly by only recording MSI data of specific areas of interest such as FIB-milled grid squares. In addition, our workflow is extremely robust and can be applied even to “old” cryo-ET samples — some of the grids we analyzed were stored for up to 12 months in liquid nitrogen after cryo-ET data acquisition, and for several weeks at 4°C after freeze-drying. Taken together, MSI can be used on any cryo-ET samples that were not discarded but remained in storage after EM data acquisition.

While MSI data is not as comprehensive as transcriptomics or proteomics, we show that it nevertheless bears sufficient discriminatory power to assign cellular species and subtypes in a fraction of the time needed for a sequencing or proteomics experiment. As introduced here, MSI largely captures different lipid species, which represents a class of molecules that has remained severely underrepresented in cryo-ET research, despite a prevelance of studies examining cellular membranes (Glushkova et al., 2025, Medina et al., 2025). In addition, derivatization approaches (which are typically used to make certain compound classes accessible in MALDI imaging) could be implemented in this workflow to increase detectability of specific analytes such as neurotransmitters or metabolites (Shariagatorji et al., 2019). This could expand the scope of molecular analyses significantly.

One clear limitation of MSI at present, however, is its limited spatial resolution. Our pixel size of 5 µm comprises state-of-the-art MSI resolution, which is sufficient for most eukaryotic cells but not capable of distinguishing most prokaryotic cells. Other custom and developmental MALDI setups offer sub-2-µm pixel sizes (Young et al., 2025, Kompauer et al., 2016, Potthoff et al., 2025, Young et al., 2025), but employ a transmission-based laser setup, which is incompatible with our current grid holder design, or have a relatively slow laser that would make scanning a full grid very time-consuming. Further development of these setups could, however, significantly improve resolution and sensitivity for on-grid MSI. Sub-micrometer resolution, such as recently shown for the first time by Young et al. (Young et al., 2025), would enable analysis of subcellular features and organelles, or to analyze m/z signal directly on lamellae. SIMS-MS Imaging based approaches offer this resolution already, but with very few exceptions are limited to inorganic molecules and metals (Tian et al., 2024; Costa et al., 2025; Uzoni et al., 2025). We anticipate that this technology, possibly in instruments combining SIMS-MS and EM, could allow for direct classification and selection of cells on grids for FIB-milling and cryo-ET analysis in the future.

One additional challenge in our experiments comprised the cell volume remaining after FIB-milling, with multiple lamellae having only a few pixels of the respective cells remaining (which in most cases was sufficient for classification). Going forward, we suggest to adjust the FIB-milling workflow in such a way that a larger section of the cell is left intact to provide an abundant source of signal and robust statistics for unambiguous identification post hoc.

Finally, in an effort to help bridge the structural and molecular data ecosystems, we developed and deposited a set of alignment and data conversion routines and examples for the multi-omics python framework *SpatialData* (Marconato et al, 2025): https://gitlab.mpcdf.mpg.de/anschwar/spatialdata_et. This enables full integration of various modalities including reflected light, fluorescence and MSI, as well as original and derived cryo-ET data such as grid and lamella images, tomograms, segmentations, particle positions and functional states — all in the same multi-scale coordinate system. High-accuracy co-registration is facilitated by the nature of the EM grid itself, as the repetitive background pattern (grids bars) provides an excellent network of evenly spaced and clearly recognizable reference points for alignment. The full set of correlated datasets can then directly be examined within a napari viewer (Sofroniew et al., 2025), enabling comprehensive evaluation and interpretation of the data. At present, comprehensive MSI and lipidomics data are already available for many well-studied model systems, offering a robust starting point for cellular characterization in such a correlative approach. Going forward, with more single-cell lipidomics MSI data becoming available, we anticipate that cellular classification could be conducted as routinely as clinical “Biotyping”, where pathogens are identified based on empirical MALDI-MS data (Mellmann et al., 2008). Full integration of these pipelines in the same software environment thus facilitates this emerging integration of molecular and structural data going forward.

## Conclusion

Taken together, here we present the first fully integrated pipeline using MSI for the identification and classification of cells imaged using cryo-ET. Our workflow offers fast and reliable identification of cell species and type, does not require biochemical or genetic modification, and can be used on any cellular cryo-ET sample, even after extended storage. We anticipate that integration of MSI will provide complementary and essential information for *in situ* cryo-ET, in particular for studying heterogeneous samples of mixed cell populations. To fully exploit such an integrative workflow, future development will focus on improving MSI spatial resolution to sub-micrometer pixel size as well as different derivatization strategies to expand MSI detection to e.g. metabolic marker molecules or neurotransmitters. Then this integrated approach could change the way structural biologists think about and conduct *in situ* cryo-ET experiments, as sample heterogeneity and cellular identity can be resolved quickly and robustly post hoc, providing detailed information on cell species, type, and functional state.

## Acknowledgements

We would like to thank the MPI for Biophysics, in particular Dr. Martin Beck and Dr. Werner Kühlbrandt for allowing us to access their cryo-EM infrastructure and the Frankfurt Central Electron Microscopy facility, in particular Dr. Maarten Tujtel and Mark Linder for training and continued support, as well as detailed discussions on the freeze-drying process. The Max Planck Computing and Data Facility is acknowledged for setup and maintenance of computational resources. For excellent technical assistance, we would like to thank Imke Wüllenweber (lipidomics sample preparation), Fabian Baur (grid holder production) and Steffen Rebenich (laser milling).

A.S. acknowledges funding by EMBO (postdoctoral fellowship, ALT 836-2020) and the Peter und Traudl Engelhorn Stiftung. This paper was typeset with the bioRxiv word template by @Chrelli: www.github.com/chrelli/bioRxiv-word-template

## Materials and Methods

### Cell culturing and grid preparation

#### HeLa cells

EM grids (Quantifoil Au200 R2/2) were coated with fibronectin from human plasma (0.01%) overnight. Subsequently, grids were washed with ddH_2_O and stored in ddH_2_O until use. For cell seeding, each chamber of a Millicell EZ 8-well slide was filled with 40 µl DMEM (high glucose) medium (Gibco) before transferring one grid each into each well. Cells were seeded by adding 40 µl of cell suspension to each grid (final cell count: 160,000 cells/ml). Cells were incubated overnight at 37°C, 5% CO_2_. After 24 h, grids were blotted for 4 s with Whatman® filter paper grade 1, and plunge frozen in liquid ethane using a Leica EM GP2 plunge freezer, operated at 37°C, 70% humidity.

#### Polytomella

*Polytomella* cells were cultured as described previously (Dietrich et al, 2024). In short, 50 ml MAP medium (30 mM sodium acetate, 35 mM MES, 1 mM potassium phosphate (pH 7.4), 7.4 mM NH_4_Cl, 0.3 mM CaCl_2_, 0.5 mM MgSO_4_, 1.39 µM ZnSO_4_, 0.8 µM H_3_BO_3_, 2.65 mM MnSO4, 0.74 µM FeCl_3_, 0.16 mM CuSO_4_, 0.83 µM NaMoO_4_, and 0.6 µM KI, 20 µg/L thiamine, 1 µg/L cyanocobalamin) was inoculated with 10 µl of *Polytomella* cell culture and grown at room temperature without agitation for 48 h until OD_600_ = 0.9. For grid preparation, 3 µl culture was applied to EM grids (Quantifoil Cu200 R2/2) prior to blotting (3.5 s with Whatman® filter paper grade 1) and plunge freezing in liquid ethane using the Leica GP2 plunge freezer, operated at 25°C, 70% humidity.

#### HeLa & Polytomella species mixing experiment

HeLa cells were seeded as described above. In addition, prior to freezing, 3 µl of *Polytomella* cell culture (OD_600_ = 0.9) were applied to the grid, blotted for 4 s with Whatman® filter paper grade 1, and plunge frozen in liquid ethane using a Leica EM GP2 plunge freezer, operated at 37°C, 70% humidity.

#### Rat hippocampal primary brain cells

The grids containing primary rat hippocampal brain cells had previously been grown, acquired, and remained in liquid nitrogen storage (manuscript submitted). In this work, we used one of these grids. Briefly, rat hippocampal neurons were prepared from P0-P1 rat pups (Sprague Dawley, IGS, Crl:CD(SD), Charles River Laboratories). The hippocampus was dissected, digested with Papain (Sigma), and plated at 140,000 cells/ml on glass bottom dishes (P35G-1.5-14-C, Mattek) containing one EM grid each (Quantifoil Au200 R2/2), coated with 0.1 mg/ml poly-D-lysine (BD). After 4 h, 700 μl glia-conditioned medium was added and cells were cultured for 21 days at 37 °C in 5% CO2 with the addition of 700 μl normal growth medium (NGM; Neurobasal A (Invitrogen/Gibco) without phenol red, supplemented with B27 (Invitrogen/Gibco) and GlutaMAX (Gibco)) every week. Neurons were imaged in E4 buffer (120 mM NaCl, 3 mM KCl, 10 mM D-Glucose, 10 mM HEPES (1 M solution, Gibco 15630-056), pH adjusted to 7.4 with 1 M NaOH, supplemented with 2 mM MgSO4 and 2 mM CaCl2 on the day of the experiment) supplemented with NeuO (0.25 μM, Stemcell Technologies) and 20 mM HEPES on a Zeiss LSM 880 Examiner upright laser scanning confocal microscope, operated at 37°C, with a 20x/0.5NA water dipping objective (Zeiss W N-Achroplan 20x/0.5 M27). A 488 nm excitation wavelength was used and emitted fluorescence as well as transmitted light were collected as a 8 x 8 tile-scan with 13 z planes at a pixel size of 415 nm x 415 nm and 3.71 μm optical sections, stitched and maximum z-projected in the Zeiss ZEN software for correlation. Grids were then blotted for 10 s with Whatman® filter paper grade 1, and plunge frozen in liquid ethane using a Leica EM GP2 plunge freezer, operated at 37°C, 70% humidity.

### Cryo-FIB milling

Cryo-FIB milling was performed using a Thermo Scientific Aquilos 2 dual beam microscope. Grids were coated with organometallic platinum for 13 s using the gas injection system (GIS) and sputter coated with inorganic platinum for 20 s. At this point, the previously acquired NeuO images were imported and correlated in 2D using the MAPS 3 software with three landmark points. FIB sample thinning was conducted manually at a milling angle of 11° in several steps: First, vertical micro-expansion joints were cut at 1 nA current, followed by horizontal cuts spaced 5 μm apart at 1 nA, 3 μm cuts at 0.5 nA, 1 μm cuts at 0.3 nA, and finally lamella polishing at 180 - 230 nm distance and 30 pA current. A final sputter coating of inorganic platinum for 1-2 s was performed and grids were stored in liquid nitrogen until TEM acquisition. For mixed HeLa / *Polytomella* grids, lamellae were prepared automatically using the AutoTEM software.

### Cryo electron tomography

TEM grid/lamella overviews as well as tilt series were previously acquired (manuscript submitted). Briefly, all data used in this study was acquired via SerialEM (Mastronarde, 2003) on a Titan Krios G2 transmission electron microscope (Thermo Scientific), operated at 300 keV in EFTEM mode equipped with a BioQuantum K3 imaging filter (Gatan). In order to first identify suitable lamellae regions and then suitable subcellular regions, lower magnification montage images were acquired at 135x / 6,500x nominal magnification and 0 / -50 μm defocus respectively. Suitable positions for tilt series acquisition were then acquired using the tilt controller via the dose symmetric tilt scheme (Hagen et al, 2017) from -49° to +63° (with +11° pre-tilt) in 2° increments with a pixel size of 2.682 Å/px (33,000x nominal magnification) and an energy filter slit width of 20 eV. A total dose of ∼120 e-/Å2 was distributed over the corresponding 58 tilts, amounting to ∼2.2 e-/Å2/tilt respectively and divided into 10 frames with on-the-fly motion correction in SerialEM. Successive tilt series were acquired at -2.5 to -4.5 μm defocus with - 0.25 μm increment. Settings were identical for all datasets.

#### Subtomogram averaging of neuronal ribosomes

The full dataset containing ∼284,000 ribosomes was previously processed to a global resolution of the consensus map of 5.4 Å (manuscript submitted). For this work, this dataset was subset to only contain ribosomes originating from the single grid depicted in **Figure 4A**. This yielded 15,451 particles. A particle list containing these was subjected to automatic 3D refinement in region (v3.1) (Zivanov et al., 2018), followed by three rounds of refinement in M (v1.0.9) (Tegunov et al., 2021), with one temporal pose and three sub-iterations, solving for particle poses and 1 x 1 image warp grid for two rounds and 2 x 2 image warp grid for the final round. The final global (masked) resolution was 6.3 Å. Translation state designations depicted in **Figure 5** were adapted from the classification of the full dataset.

#### HeLa/*Polytomella* tomogram reconstruction and membrane segmentation

Acquired tilt series for mixed HeLa/*Polytomella* grids were reconstructed using AreTomo2 (Zheng et al., 2022) with following flags: ‘AreTomo2 - InMrc ../${tomoNumber}.mrc -OutMrc ${tomoNumber}.mrc -LogFile 1 - AngFile ${tomoNumber}.rawtlt -VolZ 2000 -PixSize 2.682 -Kv 300 – Cs 2.7 -TiltCor 0 -OutBin 6 -Wbp 1 -DarkTol 0.7 -OutImod 2 -FlipVol 1’ and membranes were automatically segmented using membrain-seg (v1, model 10?) (Lamm et al., 2022). The resulting segmentations were cleaned and selected for mitochondrial membranes in ChimeraX (v1.9) (Pettersen et al., 2021) using ‘erase sphere’ and ‘hide dust’.

#### Freeze-drying and storage

For freeze-drying, cryo-EM grids were loaded (back) into the Thermo Scientific Aquilos 2 dual beam microscope. At a constant pressure of ∼2-4 ×10^-7^ mbar, the stage temperature was gradually raised to ∼-90°C - -85°C by adjusting the nitrogen flow rate via the built-in electronic integrated nitrogen gas flow controller (Bronkhorst FlowView software). This target temperature was chosen to maximize sublimation rate while minimizing re-crystallization, but depending on the application, a higher temperature and thus faster drying processes might be preferable. The exact flow rate necessary depends on the microscope setup and will have to be determined empirically. Slight changes to hardware configuration (e.g. exchange of tubing) have led to changes in target flow rate on our setup in the past. Cells and lamellae were monitored before and during the drying process via an integrated iFLM widefield fluorescence microscope in reflected light mode and judged by a change in the interference patterns over time (**Figure S1, Movie S1**). No imaging via SEM or FIB beams was performed once the temperature was raised to avoid radiation damage. Drying generally occurred in two phases: First the surrounding medium disappeared, followed by specimen shrinkage. Once no more changes were observed (typically on the order of 4 hours), the microscope was brought to room temperature by reducing the flow to 0, typically overnight. Grids were then removed from the microscope and stored in a desiccated environment at 4°C until further use. We use a 50 ml plastic tube with silica beads in a refrigerator.

#### Reflected light imaging

Reflected light images of freeze-dried grids form the basis for mask generation and provide ultrastructural information for the interpretation of other modalities in several places throughout this manuscript. In order to acquire these, freeze-dried (clipped) auto grids were deposited cells facing downwards into glass bottom dishes (P35G-1.5-14-C, Mattek) and images were acquired in reflected light mode on an inverted Zeiss LSM 980 laser scanning confocal microscope using a 20x air objective (Plan-Apochromat 20x/0,8 M27, 420650-9903-000) and 488 nm excitation light as circular tile scan with several z sections, stitched and maximum z projected on the fly using default settings in Zeiss Zen.

#### MSI autogrid holder design & fabrication

The MSI autogrid holder was designed to securely fix EM grids during matrix applications and imaging measurements. It is compatible with standard target frames for specimen carriers and features precisely milled grooves that ensure stable x/y-positioning of the grids while compensating for the sample height caused by clamping rings. Fixation in the z-direction is achieved by flat springs attached to magnets integrated into the adapter base. The design was created using the CAD software *Solidworks 2022* (Dassault Systèmes). The base holder was manufactured on a 5-axis CNC machining center (HERMLE C 20 U, iTNC 530 Heidenhain control). The springs were produced using a laser processing center. A corresponding CAD file has been uploaded to: gitlab.mpcdf.mpg.de/anschwar/spatialdata_et/-/tree/main/raw_data

#### MSI matrix application by sublimation

Matrix application was performed using a glass sublimation apparaturs (Ace Glass) that was equipped with a custom-built matrix reservoir insert at the bottom. The EM-grids were mounted in the holder and the whole assembly attached to the cooling finger of the apparatus. Subsequently 25 mg DHB matrix were dissolved in acetone and distributed evenly in the pre-heated reservoir of the sublimation chamber . After the evaporation of solvent the chamber was closed and the sublimation was carried out for 1.5 min at 170 °C and a vacuum of 5×10^-2^ mbar.

#### MSI data acquisition

MSI data were acquired on a timsTOF fleX MALDI-2 instrument (Bruker Daltonics) equipped with microGRID. The instrument was controlled via timsControl 6.1.4 and flexImaging 7.6. Measurements were performed in positive mode with a mass range from 300-1700 m/z and TIMS enabled with a mobility range from 0.7-1.95 V×s/cm^2^. Mass calibration was performed with red phosphorus clusters on the target and ion mobility was calibrated with ESI-L Low Concentration Tuning Mix (Agilent Technologies). All imaging experiments were acquired with a raster size of 5 × 5 µm and 30 laser shots/pixel. The MALDI-2 post-ionization laser was kept deactivated throughout this study.

#### MSI data processing and compound annotation

MSI data files were imported into SCiLS Lab and feature finding performed with the T-ReX^3^ algorithm using 10 ppm mass and 2 Å mobility interval widths, a 4 × 4 neighborhood size, 100% spectra coverage and a minimum intensity of 50. All features were then exported as ome.tif image files for further spatial data analysis.

Compound annotations were obtained by matching exact masses and mobilities against target lists and rule-based annotations in Meta-boScape 2024 and mzmine 4.7. Where possible, lipid annotations were further validated with on-grid MS/MS experiments or LC-MS based lipidomics data from bulk cell extracts.

#### Data processing

##### Cross-correlation and clustering to identify cell-specific m/z channels

All identified peaks were exported as .ome.tif image files in the SCiLS Lab software without normalization and hotspot removal, and imported for analysis in python via the ‘tifffile’ library. In order to identify channels with characteristic localization to cells, grid bars, carbon film etc., we computed a cross-correlation matrix for all normalized channels (e.g. **Figure 2A**), using ‘numpy.corrcoef()’ with ‘sklearn.preprocessing.StandardScaler’. This cross-correlation matrix was segmented via hierarchical clustering (‘scipy.cluster.hierarchy.linkage()’ and ‘scipy.cluster.hierarchy.fcluster()’). In order to identify clusters characteristic of cells, pre-selected marker channels encoding for several phosphatidyl choline species were used. All clusters containing such marker channels were then extracted and subjected to a second round of crosscorrelation and clustering to identify homogeneous vs heterogeneous channels (e.g. **Figure 2B**). All corresponding code including analysis notebooks can be found at: https://gitlab.mpcdf.mpg.de/anschwar/spatialdata_et

##### Registration of imaging modalities

For precise registration of the various imaging modalities (fluorescence light microscopy, reflected light microscopy, transmission and scanning electron microscopy, mass spectrometry imaging), an identical set of landmark points was chosen for each image. The landmarks were placed in the same order and orientation for all modalities. If the full grid was visible the first point was chosen on the center grid intersection, which has a unique asymmetric shape for Quantifoil grids. Five points were then chosen going in each cardinal direction starting with the long, round side, followed by the short round side, the sharp edge and no edge. This order is arbitrary but has to be kept consistent between grids to account for potential flipping between imaging modalities. The landmark points were saved and imported into each software package used subsequently. For quantification, registration was then performed using the “Landmark correspondences” (https://github.com/axtimwalde/mpicbg/blob/master/mpicbg_/src/main/java/Transform_Roi.java) plugin in FIJI (Schindelin et al., 2012). MSI landmark points were taken as source and RL landmark points as template, with “affine” transformation class, “least squares” and no interpolation. Target registration error estimation was performed using the ec-CLEM (Paul-Gilloteaux et al., 2017) plugin in icy (de Chaumont et al., 2012), and for visualization, the spatialdata (Marconato et al., 2025) library in python and napari (Sofroniew et al., 2025) was used, each as described below.

##### Creation of individual cell masks

Step by step illustrations for the entire workflow are available in **Figure S4A** exemplified for the HeLa grid displayed in **Figure 2E**. First, individual cells were manually segmented on the reflected light image using the “labels” function in napari and saved as a binary image. In parallel, a mask covering the acquired MSI area was produced via maximum intensity projection of all m/z channels followed by the “Fill holes” operation in FIJI. This binary mask was then registered and upsampled to the reflected light coordinate system using a set of landmark points as described above using “Landmark correspondences” in FIJI. The resulting MSI area mask and the cell mask were then multiplied and individual cells were segmented using the watershed implementation in FIJI with manual curation.

##### Creation of soma / neurite masks

For the neuron grid displayed in **Figure 4A**, step by step illustrations for the entire workflow are available in **Figure S6E**. Individual cell somata and neurites were automatically segmented in the NeuO fluorescence microscopy image using the “Pixel classification” workflow in ilastik (Berg et al., 2019) using four labels: Cell somata, neurites, grid background, and empty regions. To account for defects during MSI preparation, a defect mask was produced from the reflected light image using “MinError(I)” thresholding in FIJI. This defect mask was registered to the cell mask as described above and multiplied to create a defect-corrected cell mask. Finally, to account for the MSI acquisition area, an MSI area mask was created as above via maximum intensity projection of all m/z channels followed by “Fill holes” in FIJI. This mask was also registered, transformed and upscaled into the fluorescence microscopy coordinate system and multiplied with the defect-corrected cell mask to produce the final mask used for quantification of m/z intensities in somata vs neurites described below.

##### Extracting MSI intensities for individual cell ROIs

In order to now extract intensity values for each ROI and each channel, the newly created mask alongside the reflected light and MSI image stacks were opened in FIJI. Registration points were manually chosen along the grid pattern in reflected light, transmission light or in grid bar specific m/z channels (**Figure S6**). Using a FIJI macro, for each channel, that channel was duplicated, registration positions were applied using the ROI manager and the channel was transformed to be registered with the mask using the “Landmark correspondences” plugin. MSI landmark points were taken as source and RL landmark points as template, with “affine” transformation class, “least squares” and no interpolation. For each ROI in the cell mask (which are now overlapping with the newly transformed MSI channel image), the mean MSI intensity values per area were then recorded using the “Measure” command in FIJI, concatenated and saved as csv file. Recorded mean m/z per area were next read and converted to a ‘pandas’ DataFrame in python. Spectra for individual m/z channels were plotted as ‘matplotlib’ bar plots for selected cells, and a principal component analysis was performed for all cells across all cell-specific channels and displayed as ‘matplotlib’ scatter plot, color-coded by the intensity of channel 744.48 m/z.

For neuronal soma/neurite ratio measurements, the same macro was used but rather than recording a mean intensity value per cell ROI, a single value for the entire neurite mask area and the entire soma mask area was recorded. These were divided and plotted as ‘seaborn’ line plot in python.

##### Target registration error estimation

In order to estimate the target registration error between MSI data and other registered modalities (such as reflected light), we used the ecCLEM (Paul-Gilloteaux et al., 2017) plugin in icy (de Chaumont et al., 2012). For this we imported both images into ec-CLEM alongside the previously selected registration points (landmarks). The transformation was applied and the local target registration error for each pixel was calculated. The resulting heatmap was then cropped to contain only the source region of MSI data acquisition and displayed using viridis.

