## Supplementary Information for "Correlative MS Imaging for *in situ* cryo-ET"

bioRxiv

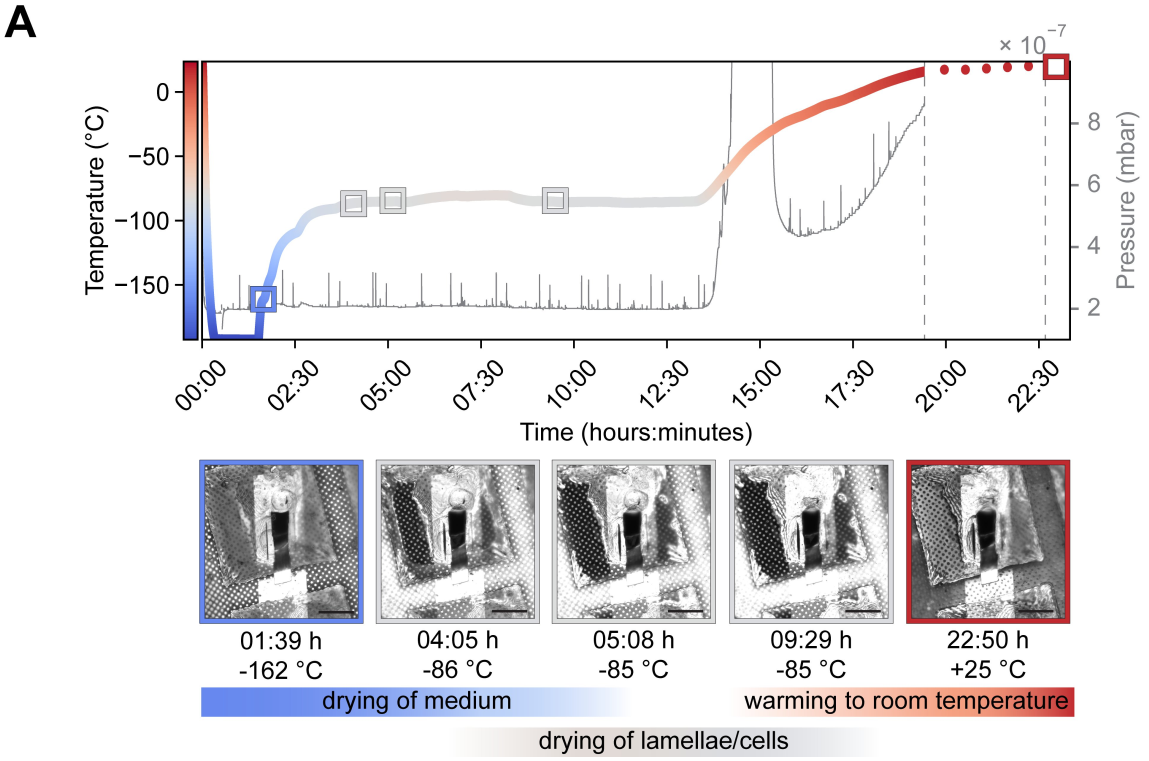


**Figure S1: Freeze-drying progression and temperature profile. (A)** Cryo-EM grids containing HeLa cells are freeze-dried within a FIB-SEM dual beam microscope chamber. Under constant chamber pressure (~$2\times10^{-7}$ mbar) grids are warmed to ~-85°C and monitored using reflected light imaging via a built-in widefield microscope. Individual snapshots are displayed showing the drying of residual cell culture medium, followed by drying of cells and lamellae, as visible by interference patterns. When no further drying is observed, the stage is warmed to room temperature. The final 3 h of drying were not recorded and are instead interpolated.

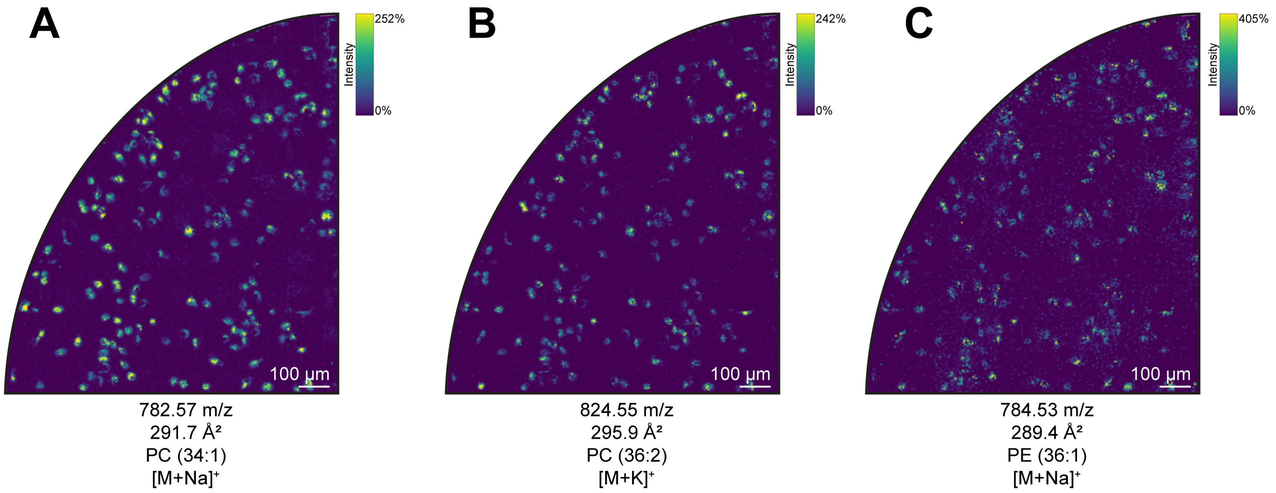


**Figure S2: Example images for HeLa cell-specific lipids. (A)** PC 34:1 (m/z 782.56, [M+Na]+), **(B)** PC 36:2 (m/z 824.55, [M+K]+), **(C)** PE 36:1 (m/z 784.53 [M+Na]+).

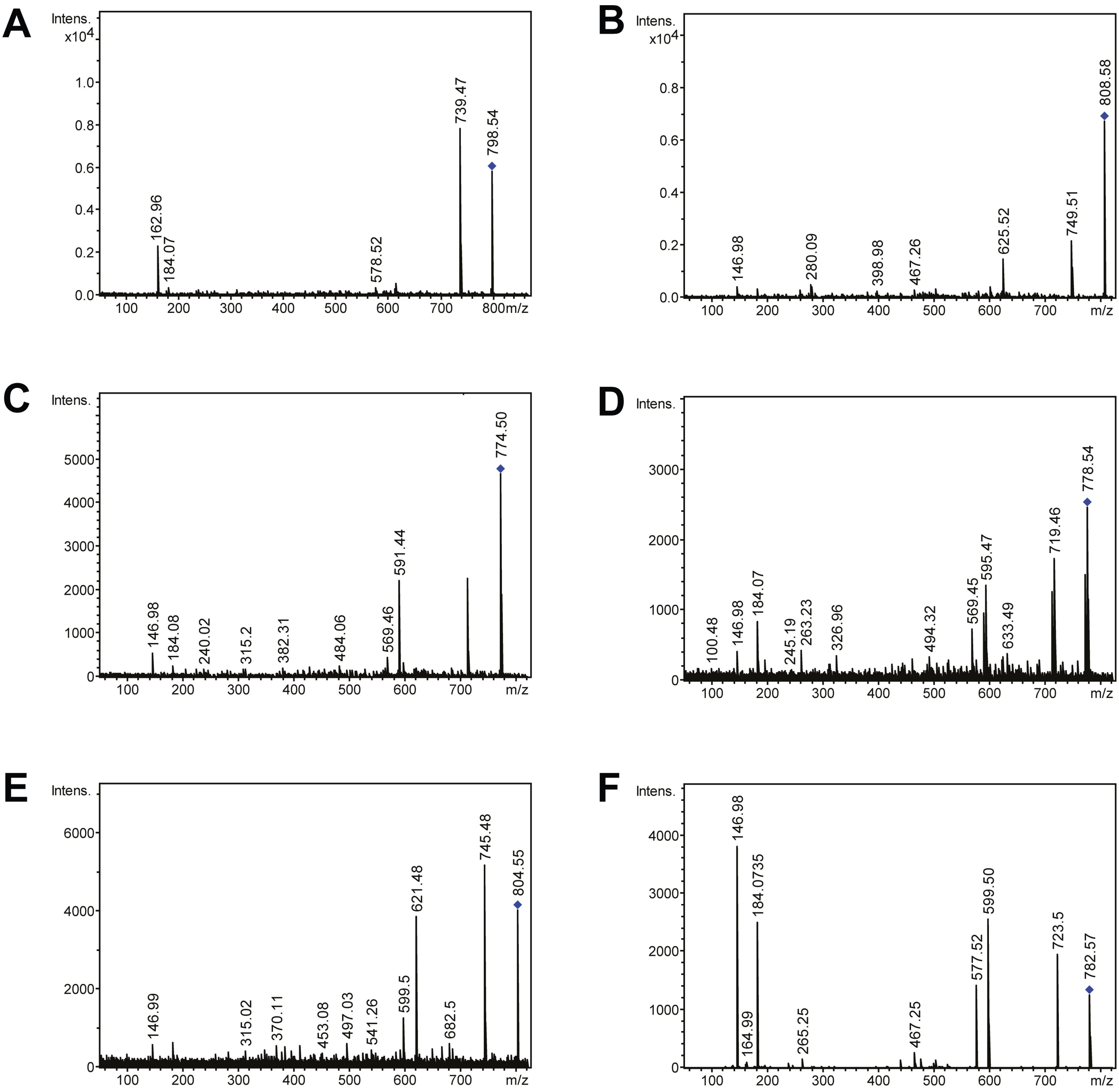


**Figure S3: On-grid MS/MS spectra of various grids. (A)** m/z 798.54 PC 34:1 (HeLa), **(B)** 808.58 PC 38:5 (HeLa), **(C)** 774.5 PC 34:5 (*Polytomella*), **(D)** 778.54 PC 34:3 (*Polytomella*), **(E)** 804.55 PC 36:4 (Neurons), **(F)** 782.57 PC 34:1(Neurons). Precursor ions are marked by a blue diamond.

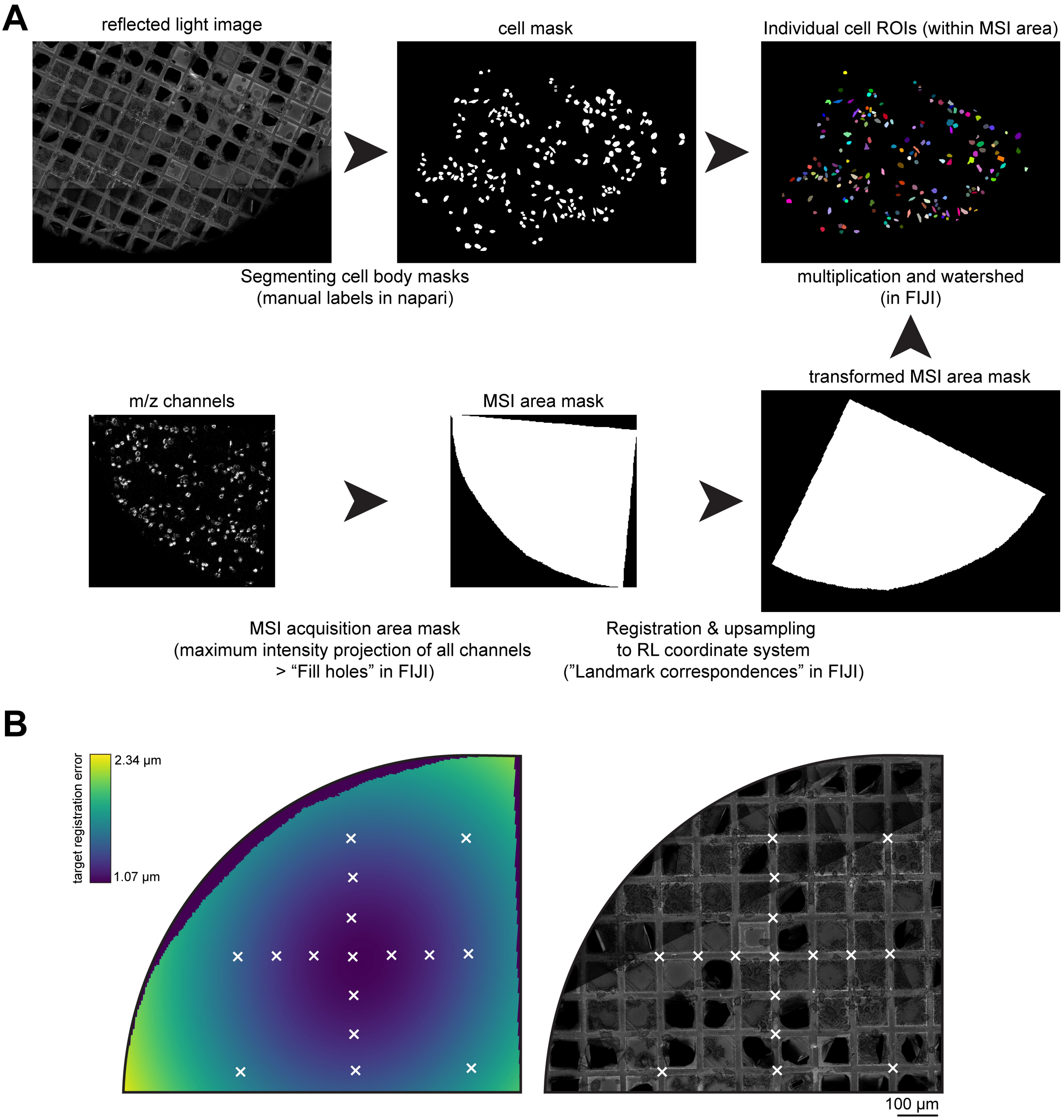


**Figure S4: HeLa grid cell mask creation and registration error estimation. (A)** Cell mask generation for HeLa grid. Individual cell masks were manually created from RL image. A MSI area mask was created via maximum intensity projection of all MSI channels and registered with landmark points displayed in (A). Images were multiplied and single cell masks generated via watershed. **(B)** Target registration error estimation as calculated for mass spectrometry imaging (MSI) to reflected light (RL) registration based on displayed landmark points, color-coded in viridis. Pixel size of MSI = 5 µm, reflected light = 0.33 um.

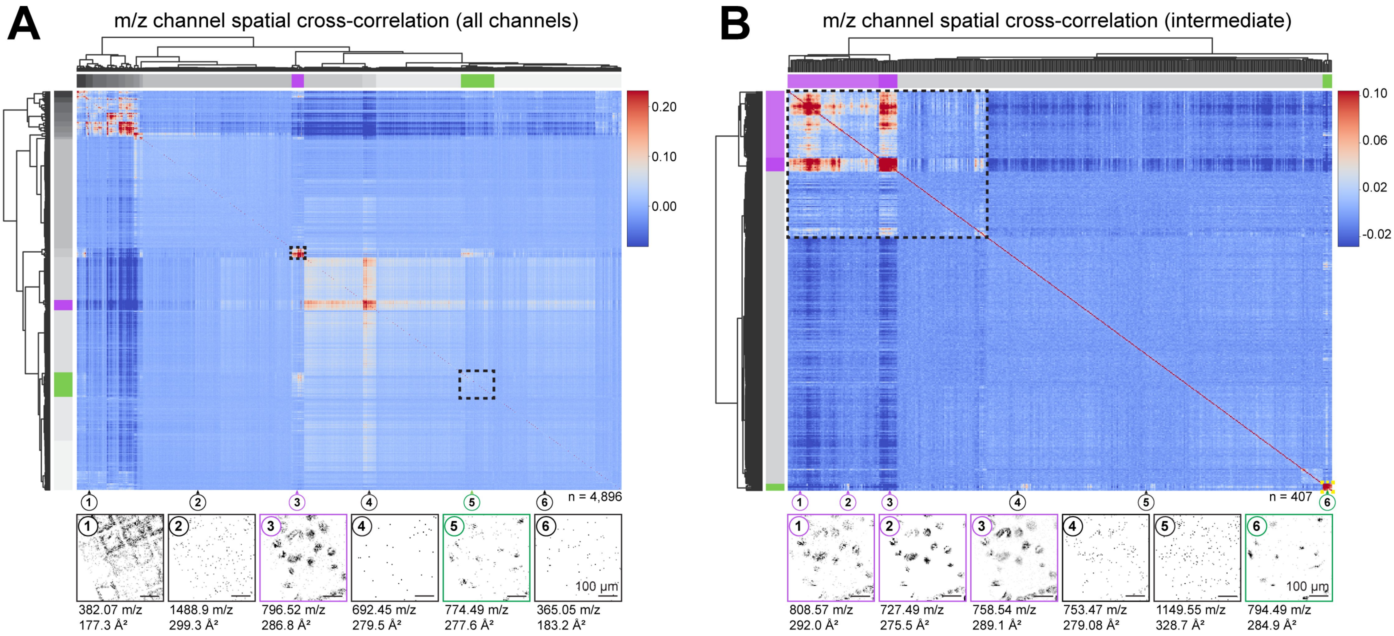


**Figure S5: Pre-filtering of cell-specific m/z channels via hierarchical clustering of normalized spatial correlations of mixed HeLa/*Polytomella* grid. (A)** Level 1: All m/z channels (n=4,896). Clusters containing cell-specific channels are highlighted in green (*Polytomella*) and magenta (HeLa). Example m/z channels for six selected clusters are shown (bottom panels). **(B)** Level 2: Second round of clustering with extracted m/z channels from (A). Channels with poor signal-to-noise ratio are removed, highlighted clusters are designated as cell-specific and used in **Figure 3A** (n=407).

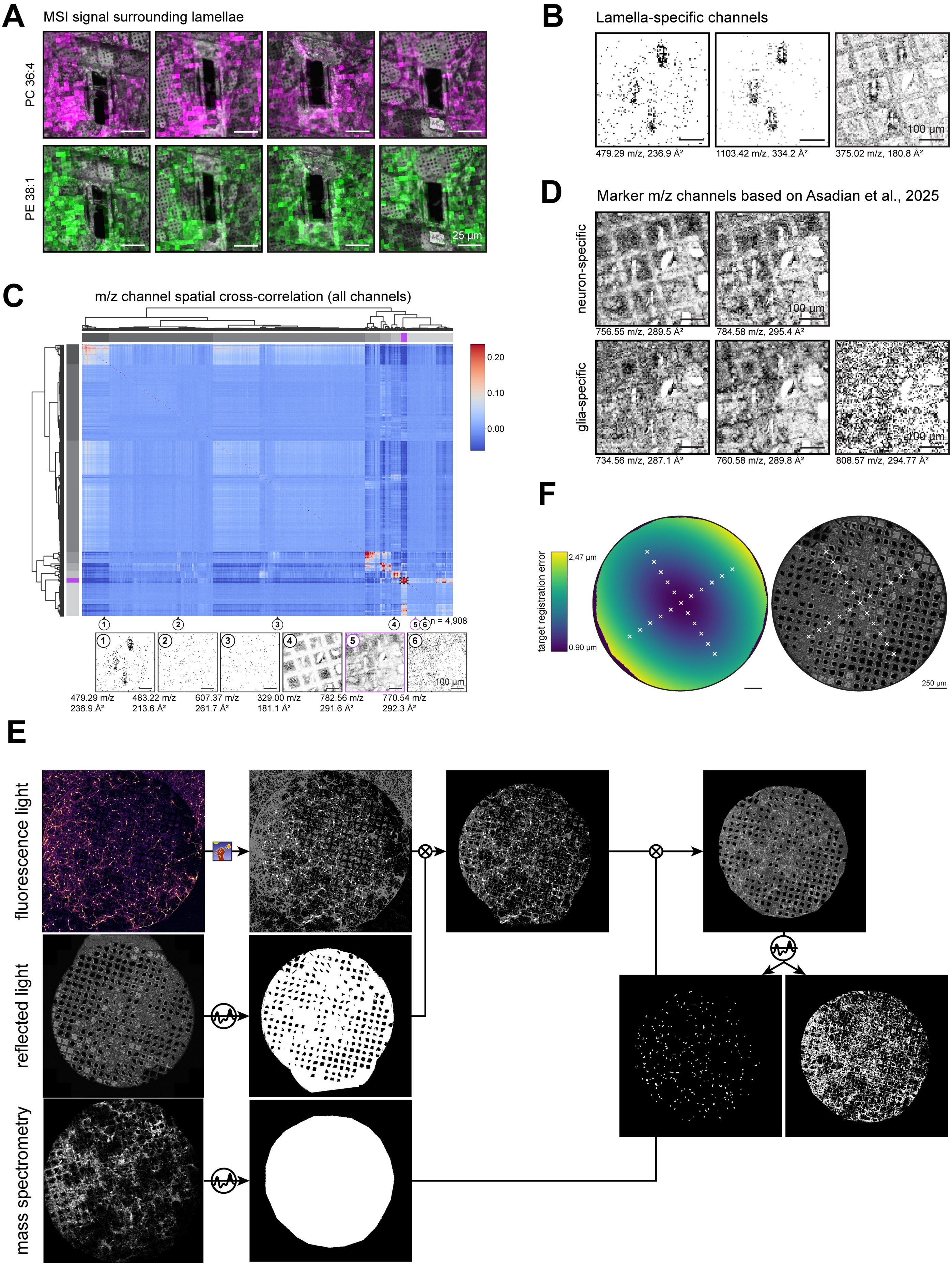
A
MSI signal surrounding lamellae
PC 36:4
PE 38:1
25 µm
B
Lamella-specific channels
479.29 m/z, 236.9 Å²
1103.42 m/z, 334.2 Å²
375.02 m/z, 180.8 Å²
100 µm
D
Marker m/z channels based on Asadian et al., 2025
neuron-specific
756.55 m/z, 289.5 Å²
784.58 m/z, 295.4 Å²
glia-specific
734.56 m/z, 287.1 Å²
760.58 m/z, 289.8 Å²
808.57 m/z, 294.77 Å²
100 µm
C
m/z channel spatial cross-correlation (all channels)
0.20
0.10
0.00
n = 4,908
479.29 m/z
236.9 Å²
483.22 m/z
213.6 Å²
607.37 m/z
261.7 Å²
329.00 m/z
181.1 Å²
782.56 m/z
291.6 Å²
770.54 m/z
292.3 Å²
100 µm
F
target registration error
2.47 µm
0.90 µm
250 µm
E
fluorescence light
reflected light
mass spectrometry

**Figure S6: Examples of lamella-enriched m/z signal and marker channels, pre-filtering of cell-specific channels, registration error estimation, and mask creation of neuron grid. (A)** MSI signal surrounding primary rat hippocampal neuron lamellae. Example m/z intensities for PC 36:1 () and PE 38:1 () are shown overlaid on top of reflected light images for four lamellae of the same grid shown in **Figure 4A**. **(B)** **(B)** Some m/z channels display increased intensity surrounding lamellae, indicating beam-induced changes in these molecular species. **(C)** Pre-filtering of cell-specific m/z channels via hierarchical clustering of normalized spatial correlations of neuron grid. All m/z channels (n=4,908). Clusters containing cell-specific channels are highlighted in magenta. Example m/z channels for selected clusters are shown (in bottom panels), highlighted clusters are designated as cell-specific and used in **Figure 4D**. **(D)** Images of neuron- and glia-enriched m/z values identified in (Asadian et al., 2025) **(E)** Soma/neurite mask generation for neuron grid. Initial soma/neurite segmentation was performed on NeuO fluorescence light image via neural network-based pixel classification. A grid defect mask was created based on the reflected light image, registered using landmark points in (F), and multiplied with the soma/neurite mask. A final mask marking the MSI acquisition area was created by maximum intensity projection of all m/z channels, registered, and also multiplied to create the final soma/neurite mask. Soma and neurite masks were applied separately for m/z intensity measurements. **(F)** Target registration error estimation as calculated for mass spectrometry imaging (MSI) to reflected light registration based on displayed landmark points, color-coded in viridis. Pixel size of MSI = 5 um, reflected light = 0.33 um.
